# Structural basis of bifunctionality of *Sinorhizobium meliloti* Clr, a cAMP and cGMP receptor protein

**DOI:** 10.1101/2022.11.04.515266

**Authors:** Laura Werel, Neda Farmani, Elizaveta Krol, Javier Serrania, Lars-Oliver Essen, Anke Becker

**Author notes:** Corresponding authors: Lars-Oliver Essen and Anke Becker. Authors contributed equally to this work (Laura Werel, Neda Farmani).

## Abstract

In bacteria, Crp-Fnr superfamily transcription factors are the most ubiquitous receptor proteins of 3’,5’-cyclic adenosine monophosphate (cAMP) and 3’,5’-cyclic guanosine monophosphate (cGMP). The prototypic *Escherichia coli* CAP protein represents the main CRP subclass and is known to bind cAMP and cGMP, but to mediate transcription activation only in its cAMP-bound state. In contrast, both cyclic nucleotides mediate transcription activation by CRP subclass G protein Clr of *Sinorhizobium meliloti*. We present crystal structures of apo-Clr, and Clr•cAMP and Clr•cGMP bound to the core motif of the palindromic Clr DNA binding site (CBS). We show that both cyclic nucleotides shift ternary Clr•cNMP•CBS-DNA complexes to almost identical active conformations. Unlike the situation known for the *E. coli* CAP•cNMP complex, in the Clr•cNMP complex, the nucleobases of cGMP and cAMP are in the *syn-* and *anti*-conformation, respectively, allowing a shift to the active conformations in both cases. Isothermal titration calorimetry measured similar affinities of cAMP and cGMP binding to Clr in presence of CBS core motif DNA (K_D_^cNMP^ 16 μM). However, different affinities were determined in absence of this DNA (K_D_^cGMP^ 24 μM; K_D_^cGMP^ 6 μM). Sequencing of Clr co-immunoprecipitated DNA as well as Electrophoretic Mobility Shift and promoter-probe assays expanded the list of experimentally proven Clr-regulated promoters and CBS. This comprehensive set of CBS features conserved nucleobases, which are in agreement with the sequence readout through interactions of Clr amino acid residues with these nucleobases, as revealed by the Clr•cNMP•CBS-DNA crystal structures.

**IMPORTANCE:** Cyclic 3’,5’-adenosine monophosphate (cAMP) and cyclic 3’,5’-guanosine monophosphate (cGMP) are both long known as important nucleotide second messenger in eukaryotes. This is also the case for cAMP in prokaryotes, whereas a signaling role for cGMP in this domain of life has been recognized only recently. Catabolite repressor proteins (CRPs) are the most ubiquitous bacterial cAMP receptor proteins. *Escherichia coli* CAP, the prototypic transcription regulator of the main CRP subclass, binds both cyclic mononucleotides, but only the CAP•cAMP complex promotes transcription activation. In contrast, CRP subclass G proteins studied so far, are activated by cGMP, or both by cAMP and cGMP. Here, we report a structural analysis of the bifunctional cAMP- and cGMP-activatable Clr from *Sinorhizobium meliloti*, how binding of cAMP and cGMP shifts Clr to its active conformation, and the structural basis of its DNA binding site specificity.

## INTRODUCTION

3’,5’-cyclic adenosine monophosphate (cAMP) is one of the most ubiquitous nucleotide second messengers in prokaryotes and eukaryotes. It was the first second messenger to be described, initially for its role in hormone-dependent signal transduction by eukaryotes (1, 2), later for catabolite repression in bacteria like *Escherichia coli* (3). Since then, a myriad of biological processes regulated by cAMP have been reported in bacteria, such as carbon metabolism, biofilm formation, type III secretion, virulence, and symbiosis (4, 5). Another cyclic mononucleotide second messenger widespread in eukaryotes is 3’,5’-cyclic guanosine monophosphate (cGMP) (6). Involvement of cGMP in bacterial regulation has been recognized only recently (7).

In bacteria, cyclic AMP receptor proteins (CRPs) are among the best characterized transcription factors and model for allosteric regulation (3, 8-10). Physiological effects of gene regulation by CRP-like proteins were characterized in various bacterial species. They are highly versatile, controlling expression of at least over 200 genes in *E. coli* and almost as many in *Mycobacterium tuberculosis* and *Corynebacterium glutamicum* (11-*15*).

CRPs belong to the superfamily of Crp-Fnr transcription regulators, which are composed of an N-terminal nucleotide binding domain and a C-terminal helix-turn-helix motif (16). These domains are each conserved in eukaryotes and prokaryotes (4, 17). Upon DNA binding, the C-terminal domain of CRPs is able to interact with the bacterial RNA polymerase (RNAP) and to modulate promoter activation (18-20). Most of the CRPs act as transcription activators, few as repressors (6). CRP-like proteins of the Crp-Fnr superfamily were further divided into the main CRP, and CRP F and G subclasses (16). So far, all structurally characterized CRPs belong to the main CRP subclass, including *E. coli* CAP (21). These proteins are activated by cAMP only (16). Only two members of the CRP G subclass have been functionally studied. CrgA from *Rhodospirillum centenum* and Clr from *Sinorhizobium meliloti*. CrgA appeared to be solely activated by cGMP with very little cAMP-binding activity, whereas Clr is distinguished from any of the other known CRP homologues by its ability to be activated by both cAMP and cGMP (22, 23). cGMP binding has been described for *E. coli* CAP as well, albeit without stimulating DNA binding (24).

The soil-dwelling α-proteobacterium *S. meliloti* is capable of fixing atmospheric nitrogen in a symbiotic relationship with leguminous plants of the genera *Medicago, Melilotus* and *Trigonella*. The symbiotic program involves root hair infections and formation of root nodules, harbouring the nitrogen-fixing bacteria (25, 26). The *S. meliloti* Rm1021 and Rm2011 genomes encode 13 CRP-Fnr-like paralogs, including the cAMP-independent FixK regulators of microoxic respiration and nitrogen-fixation genes (3), and cAMP- and cGMP-controlled Clr (27). Eight of those are members of the CRP superfamily. Bifunctionality of Clr is especially interesting given that these *S. meliloti* genomes contain an exceptionally high number of 28 putative adenylate/guanylate cyclases (AC/GCs) genes (28). Clr and the AC/GCs, CyaD1, CyaD2 and CyaK, are implicated in repression of secondary infections of *Medicago sativa* roots (5). Moreover, Clr overexpression resulted in reduced swimming motility and increased succinoglycan production, which are traits relevant at early stages of the root nodule symbiosis (23).

In our study, we present crystal structures of apo-Clr and Clr•cNMP•DNA complexes, which to our knowledge is the first report of a subclass G CRP structural analysis. Our analysis unravels the molecular basis of Clr bifunctionality. Furthermore, we expand the set of experimentally proven Clr-regulated promoters and CBS. These feature conserved nucleobases which are in accordance with the sequence readout revealed by the Clr•cNMP•CBS-DNA crystal structures.

## RESULTS

### ChIP-seq assisted screening for candidate Clr binding sites

To identify Clr-cAMP and Clr-cGMP binding sites genome-wide in the DNA of *S. meliloti* Rm2011, we used chromatin immunoprecipitation coupled with deep sequencing (ChIP-seq). For this purpose, we first generated plasmid pWBT-Clr-CF and introduced it into *S. meliloti* strain Rm2011 Δ*clr*. pWBT-Clr-CF carries IPTG-inducible *clr-3xflag*, encoding Clr C-terminally fused to a 3xFLAG-tag. *In vivo* functionality of Clr-3xFLAG was confirmed by its ability to activate plasmid-borne *egfp* reporter gene fusions to promoters of the previously identified target genes *SMc02178* and *SMb20906* (23) in Rm2011 Δ*clr* in presence of externally added cAMP (Fig. S1). For α-FLAG antibody mediated chromatin immunoprecipitation, Rm2011 Δ*clr* carrying pWBT-Clr-CF was cultivated in TY complex medium or MOPS-buffered minimal medium (MM), and supplemented with either cAMP or cGMP. Deep sequencing of DNA enriched for Clr-bound regions and control samples representing sheared DNA isolated from cell lysates prior to immunoprecipitation yielded 3.3 to 3.9 million reads per sample (Table S1A). ChIP-seq data was processed using CLC-Genomics workbench to determine peak locations and peak shape score values, which reflect enrichment of the respective regions (Table S1C-F). Consolidation of our data obtained for the four samples yielded 873 distinct ChIP-enriched genomic regions, further referred to as ChIP peaks (Table S1G). Consistent with our previous observation that both cAMP and cGMP promote Clr binding to DNA (23), 40 out of 50 top-score ChIP-enriched regions were detected in samples from cultures supplemented with either of the two cNMPs (Table S1G). Moreover, peak shape score values correlated well between cAMP and cGMP samples (correlation factors 0.92 and 0.91 in TY and MM media, respectively) (Fig.1A). Out of 417 ChIP peaks identified within intergenic regions, 160 were located between two divergently transcribed genes, 229 between two genes transcribed in the same direction and 28 between two convergently transcribed genes (Fig.1B). The remaining ChIP peaks were situated within rRNA or tRNA genes and protein coding regions (Fig. 1B). To search for Clr binding sites (CBS) in ChIP-enriched regions, we extracted 400 bp DNA sequences surrounding the peak centers. Using the FIMO online tool (29), these regions were scanned for the core Clr binding site motif GTNNCNNNNGNNAC. This core motif was derived from the consensus sequence HGTYHCNNNNGRWACA, which represents six experimentally verified CBSs (23). Sequences matching the core motif were found in 102 ChIP-enriched regions, corresponding to a total of 107 putative CBSs (Fig. 1C and Table S1H), out of which 87 were located in intergenic regions (Table S1H). A FIMO search with relaxed settings (p-value 0.001) identified additional 693 putative CBSs with one mismatch in the core motif (relaxed-consensus CBSs) in 431 ChIP-enriched regions (Table S1I). Out of these, 89 putative CBSs were found in 51 ChIP-enriched regions, which already contained a strict consensus sequence.

**FIG 1.**
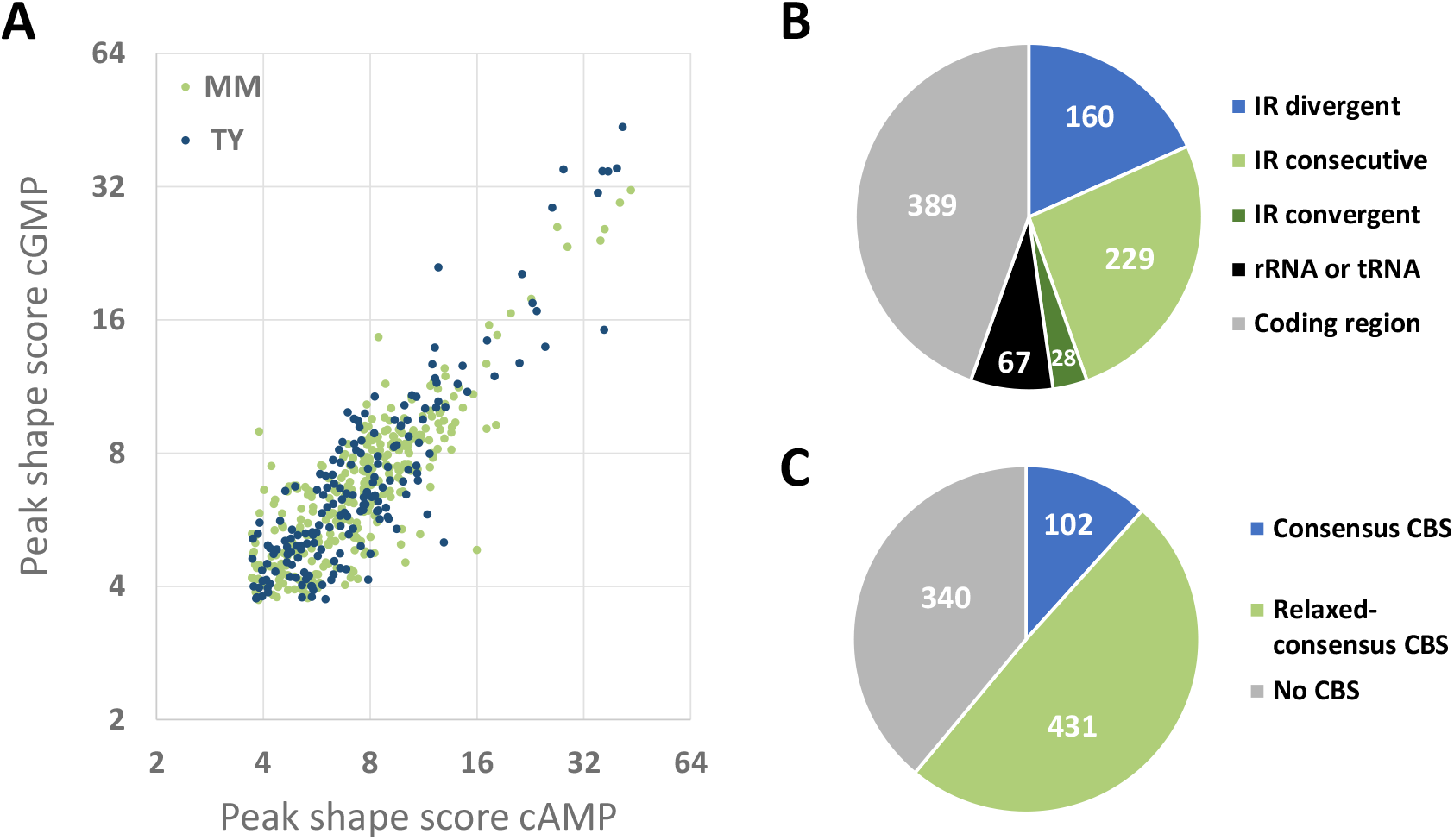
ChIP-seq-assisted screening for Clr-cAMP and Clr-cGMP binding sites. (A) Correlation between peak shape score values of ChIP-seq peaks detected in samples of cells grown in TY or MM, both supplemented with either cAMP or cGMP. (B) Location of Clr ChIP-seq peaks within genomic features. IR, intergenic region; IR divergent, IR between two genes transcribed in divergent directions; IR consecutive, IR between two genes transcribed in the same direction; IR convergent, IR between two genes transcribed in converging directions; rRNA or tRNA, rRNA or tRNA encoding region; coding region, protein coding region. (C) Presence of putative Clr binding sites (CBSs) in the vicinity of ChIP-seq peaks. Consensus CBS: GTNNCNNNNGNNAC. Relaxed consensus CBS, one mismatch to the consensus was allowed.

As expected, all previously characterized CBS sequences in the promoter regions of *SMc02178, SMc00653, SMc01136, SMc04190, cyaF2* and *SMb20906* (5, 23) were found within the ChIP-enriched regions. Among genes adjacent to ChIP peak-containing intergenic regions, 35 genes were found (Table S1J) that were previously identified in transcriptome experiments to be upregulated upon overexpression of *clr* or adenylate cyclase gene *cyaJ* (23, 30). 25 of these genes contained putative consensus and/or relaxed-consensus CBSs in their upstream non-coding regions. The remaining 10 genes lacked putative CBSs in their vicinity. Remarkably, these 10 genes were only detected as being transcriptionally activated upon *cyaJ*, but not *clr* overexpression.

### Clr-cAMP and Clr-cGMP specifically bind DNA target sequences *in vitro*

To characterize putative CBS sequences identified within ChIP-seq-enriched DNA regions and located in upstream non-coding regions, we performed electrophoretic mobility shift assays (EMSAs) for 62 genomic regions (Fig. 2A and Fig. S2). The corresponding DNA fragments that produced a complete electrophoretic mobility shift in presence of Clr and cAMP were defined as regions with a strong CBS, whereas DNA fragments that were only partially shifted were defined as containing a weak CBS (Fig. 2B). Of the 62 CBS candidates tested, 24 were classified as strong and 24 as weak CBSs, while the remaining failed to mediate a detectable band shift in the EMSAs (Fig. 2A and Fig. S2). Furthermore, Clr-cAMP produced stronger band shifts than Clr-cGMP. Not all DNA regions containing putative CBS matching the consensus produced strong band shifts, which indicates that non-conserved nucleotides within the CBS and its sequence environment might contribute to Clr-cNMP binding. For strong binding sites, we noted that the left and right halves of the palindrome tended to be enriched for pyrimidine (C/T) and purine base nucleotides (G/A), respectively (Table S1K). Conversely, putative CBSs mediating a weak or no band shift lacked a clear nucleotide preference at the non-conserved positions. Almost all DNA fragments causing a strong band shift included a CBS sequence that matches the core motif. A notable exception is the DNA fragment covering the *SMc05008* promoter region, as this region contains three overlapping putative CBSs, each with one mismatch in the core motif (Fig. 3).

**FIG 2.**
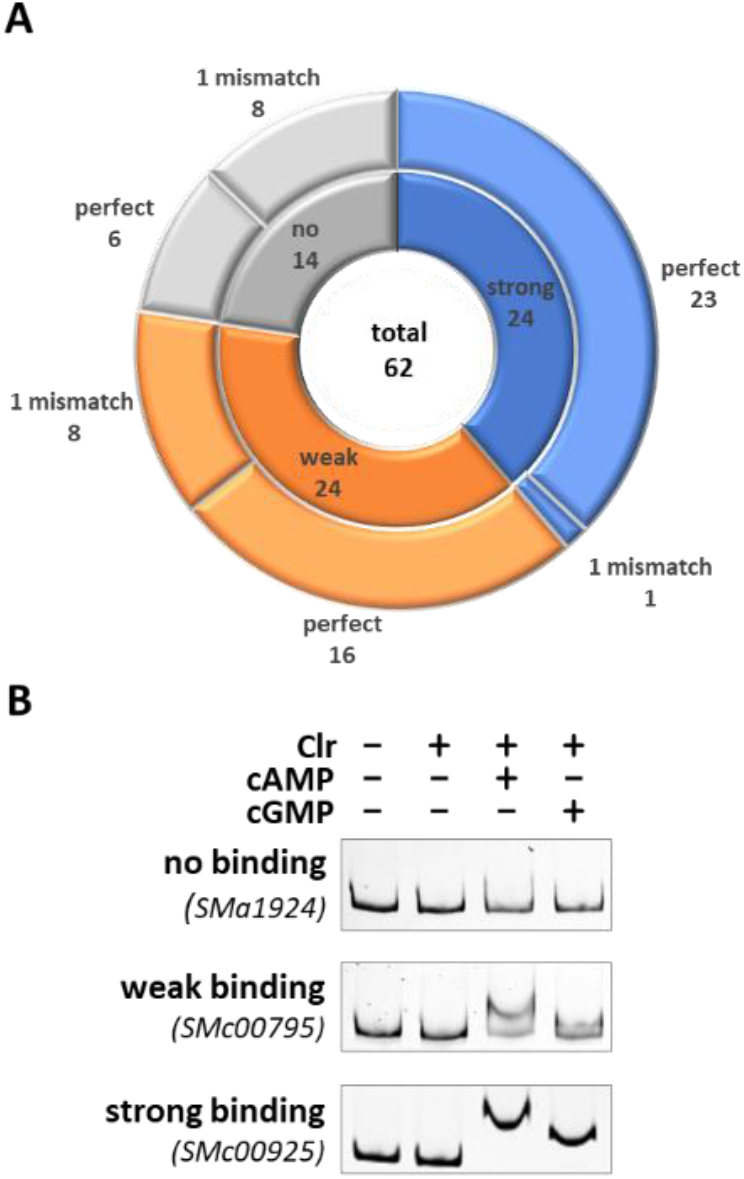
EMSA-based subclassification of CBS. All DNA fragments used for EMSAs contained at least one putative CBS matching the consensus binding site sequence either perfectly or with one mismatch. (A) EMSAs were subclassified into strong (blue), weak (orange) and no binding (grey). Numbers of perfect and imperfect binding sites are indicated for each subclass. (B) Representative EMSAs are shown for each subclass. These were performed with upstream regions of *SMa1924* (no binding), *SMc00795* (weak binding) and *SMc00925* (strong binding), which each contained a perfect match binding site.

**FIG 3.**
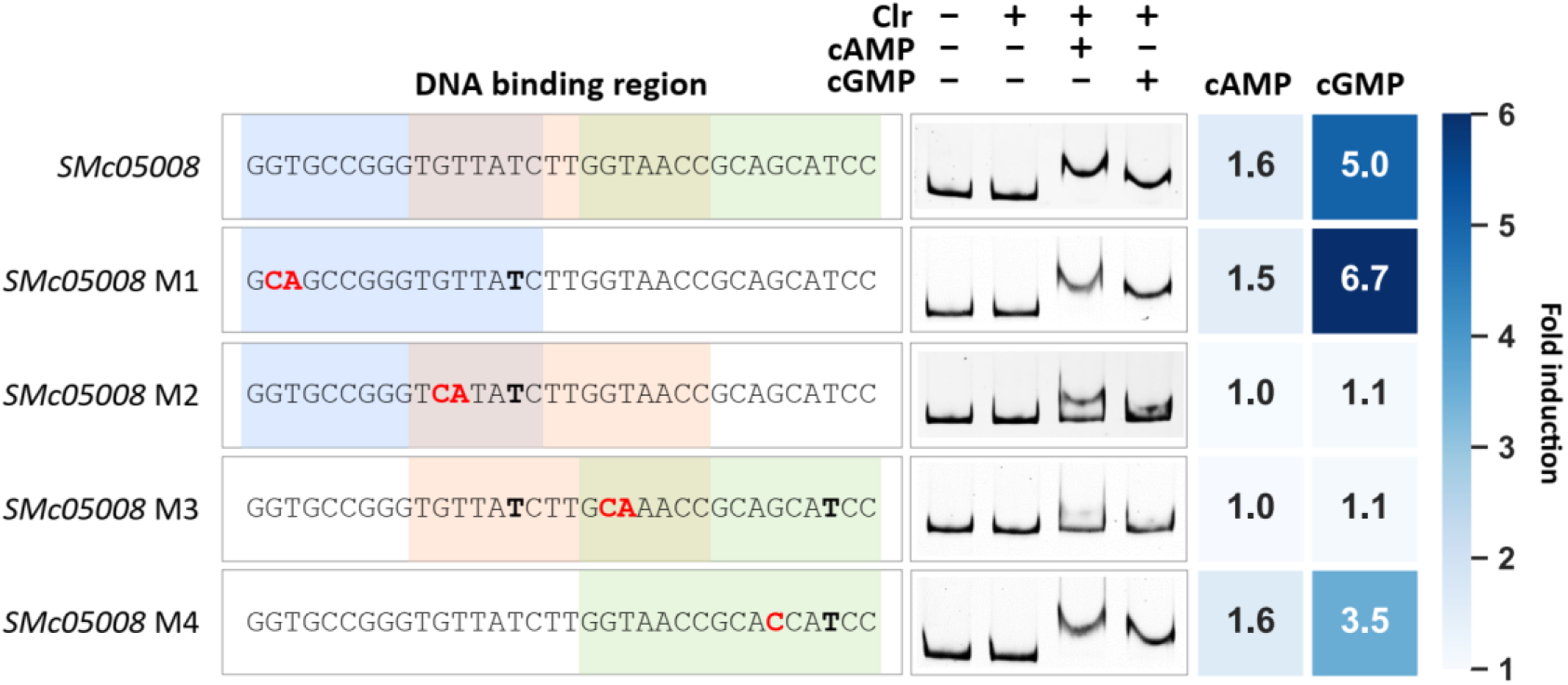
DNA binding of Clr-cAMP and Clr-cGMP at the *SMc05008* promoter. The promoter region of *SMc05008* has three possible DNA binding sites (colored), with one mismatch each. Mutations indicated in red were introduced at four different positions (M1-4). Nucleotides not matching the consensus Clr binding sequence are printed in bold. The heatmap shows the cAMP- or cGMP-induced fold change of EGFP-mediated fluorescence compared to non-inducing conditions (no cNMP added).

To verify the CBS location within DNA regions that showed strong Clr-cNMP binding, we exchanged the conserved GT in the left half of the palindrome to CA. This modification dramatically reduced Clr-cNMP binding, thus confirming the predicted CBS within these regions (Fig. S2). Out of the three overlapping putative CBS in the *SMc05008* promoter region, only mutations in conserved nucleotide positions of the overlapping putative second and third CBS inhibited Clr-cNMP binding (Fig. 3). Band shift assays with synthetic DNA fragments containing either one of the putative CBSs identified in the *SMc05008* promoter region, suggested that only the second putative CBS conferred Clr-cNMP binding (Fig. S2D). This indicates that a C→T transition at position 6 of the CBS core motif is not detrimental *per se* for Clr-cNMP binding to the DNA.

### Target promoters are regulated both by Clr-cAMP and Clr-cGMP *in vivo*

All tested CBS candidates were further investigated for Clr-cNMP dependent promoter activation *in vivo*. In case the CBS was located in non-coding regions between two opposingly transcribed genes, both putative promoter regions were tested. Plasmid-borne copies of promoter regions followed by up to 75 bp of the downstream coding regions or non-coding RNA genes fused to *egfp* were introduced to *S. meliloti* or *E. coli* for promoter activity assays.

Reporter construct-mediated fluorescence was determined in *S. meliloti* Rm2011 wild type and Rm2011 Δ*clr* upon addition of either cAMP or cGMP (Fig. 4A, Fig. S2E and Table S2). This analysis confirmed Clr-cNMP-dependent activation of 20 promoters by at least 1.5-fold either with cAMP or cGMP added. These included only promoter regions that showed a strong band shift in the EMSAs. As expected, GT→CA mutation of the first two nucleotides of the core CBS motif abolished or substantially decreased Clr-cNMP dependent promoter activation (Fig. S2E and Table S2). Regarding the three overlapping CBS in the *SMc05008* promoter region, only nucleotide exchanges that diminished Clr-cNMP binding in the EMSA also abolished promoter activation by Clr-cNMP (Fig. 3). For the majority of CBS-containing promoter fragments tested, induction by cGMP was up to four times higher than by cAMP. In total, we identified 11 novel Clr-cNMP-activated genes in addition to nine Clr-cNMP-activated genes reported previously (23, 30). The majority of the 20 confirmed Clr-cNMP regulated genes encode hypothetical proteins and, notably, eight of the 20 genes encode small proteins of less than 100 amino acids. Ten Clr-cNMP-activated genes encode proteins with an N-terminal signal peptide. Moreover, Clr-cNMP was found to directly induce transcription of *SMelC181* encoding a small non-coding RNA of unknown function. This RNA was previously identified in an RNA-seq study (31).

**FIG 4.**
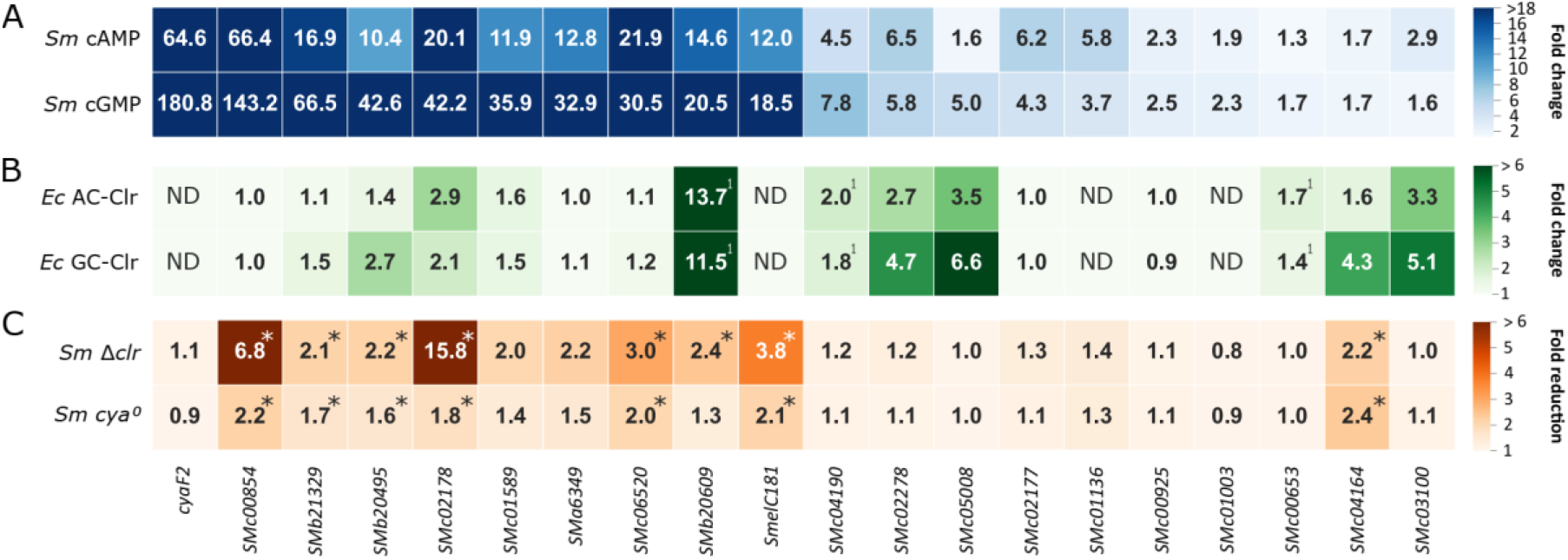
Promoter-probe measurements in different hosts. (A) Fold-change of EGFP fluorescence signal mediated by promoter-probe constructs in *S. meliloti* Rm2011 in cultures supplemented with cAMP or cGMP. Ratios derive from relative fluorescent units (RFU) of cNMP-induced against uninduced cultures. (B) Fold-change of EGFP fluorescence signal mediated by promoter-probe constructs in *E. coli* BTH101 upon induced production of an AC (CyaG1 of *S. meliloti*) or a GC (Cya2 of *Synechocystis* sp.) together with Clr (*S. meliloti*) in LB. Ratios shown derive from fluorescence units of strains under AC/GC-Clr production conditions compared to control strains producing the AC or GC but not Clr. ^1^ Values derived from (23). ND indicates data not determined. (C) Fold-reduction of EGFP fluorescence signal mediated by promoter-probe constructs in *S. meliloti* Rm2011 Δ*clr* and Rm2011 *cya*^*0*^ compared to Rm2011 wild type. *Indicate statistically significant fluorescent reduction (p < 0.05) in combination with a wild type RFU threshold of ≥ 80. Values of each biological replicate and standard deviation of samples measured in A-C are given in Tables S2-13.

Furthermore, Clr-cNMP-mediated transcription activation of overall 17 promoter regions with confirmed and putative Clr-binding motifs was analyzed in *E. coli* as a heterologous host (Fig. 4B and Table S3). We used *E. coli* strain BTH101, which is unable to produce cAMP. This strain was equipped with plasmid-borne *clr*, co-expressed with either the *S. meliloti* adenylate cyclase gene *cyaG1* or the *Synechocystis* sp. guanylate cyclase gene *cya2* under the control of an IPTG-inducible promoter. Promoter-reporter constructs were introduced on a second plasmid. We determined previously that in *E. coli* BTH101, cAMP/cGMP production alone was insufficient to activate the *S. meliloti* Clr-cNMP target promoters, and therefore *E. coli* CRP did not substitute for Clr (23). Out of the 17 promoters tested, eight were activated by Clr-cNMP in *E. coli* by at least 1.5-fold either by cAMP or cGMP production. Including the three previously described candidates *SMb20906, SMc00653* and *SMc04190* (23), a total of 11 promoters were activated by Clr-cNMP in *E. coli*. Promoters that were not clearly activated by Clr-cNMP in *S. meliloti* Rm2011 also showed no activation in *E. coli* BTH101 (Tables S2 and S3). Contrarily, five promoters (*SMc00854, SMa6349, SMc06520, SMc02177* und *SMc00925*) that were activated in *S. meliloti* Rm2011 were not activated in *E. coli* BTH101, suggesting that these promoters might require additional *S. meliloti*-specific factors for Clr-cNMP-mediated activation.

To evaluate promoter activation by Clr dependent on endogenously produced cNMPs, we compared the promoter-probe constructs conferring EGFP fluorescence between Rm2011 and its derivatives lacking either all AC/GC genes (*cya*^*0*^) or *clr* (Fig. 4C). To generate the *cya*^*0*^ strain, all 28 annotated class III AC/GC-encoding genes (28) were sequentially deleted from the Rm2011 genome. The strains were grown in medium without added cNMPs. Under these conditions, eight and seven of the 20 promoter-probe constructs mediated at least 1.5-fold (p < 0.05) reduced fluorescence levels in the Δ*clr* and *cya*^*0*^ strains, respectively, compared to the wild type (Fig. 4C and Table S4). Lack of the *clr* gene resulted in a much stronger reduction in the promoter activities of *SMc00854, SMc02178* and the sRNA gene *SmelC181* than absence of the 28 AC/GC genes.

To estimate the effect of CBS location relative to the transcription start site (TSS) on promoter activation, we gathered *S. meliloti* TSS positions, determined previously in Rm2011 and Rm1021 wild type strains (32, 33) and Rm1021 derivative RFF625c lacking all *ecf* sigma factors, and determined the CBS-TSS distances (Table S1H, I). For most promoters activated by Clr-cNMPs, CBS-TSS distances of -42 to -76 were found (Fig. 5). These promoters are thus comparable to CAP-dependent class I and class II promoters in *E. coli* (34-36). In *E. coli*, these promoters are subdivided into three classes: promoters with a CAP binding site roughly 60 to 90 bp upstream of the TSS (class I), close to the -35 element (class II), and at position -91 or further upstream (class III); the latter requiring additional binding of another transcription regulator closer to the promoter region for activation (34-36).

**FIG 5.**
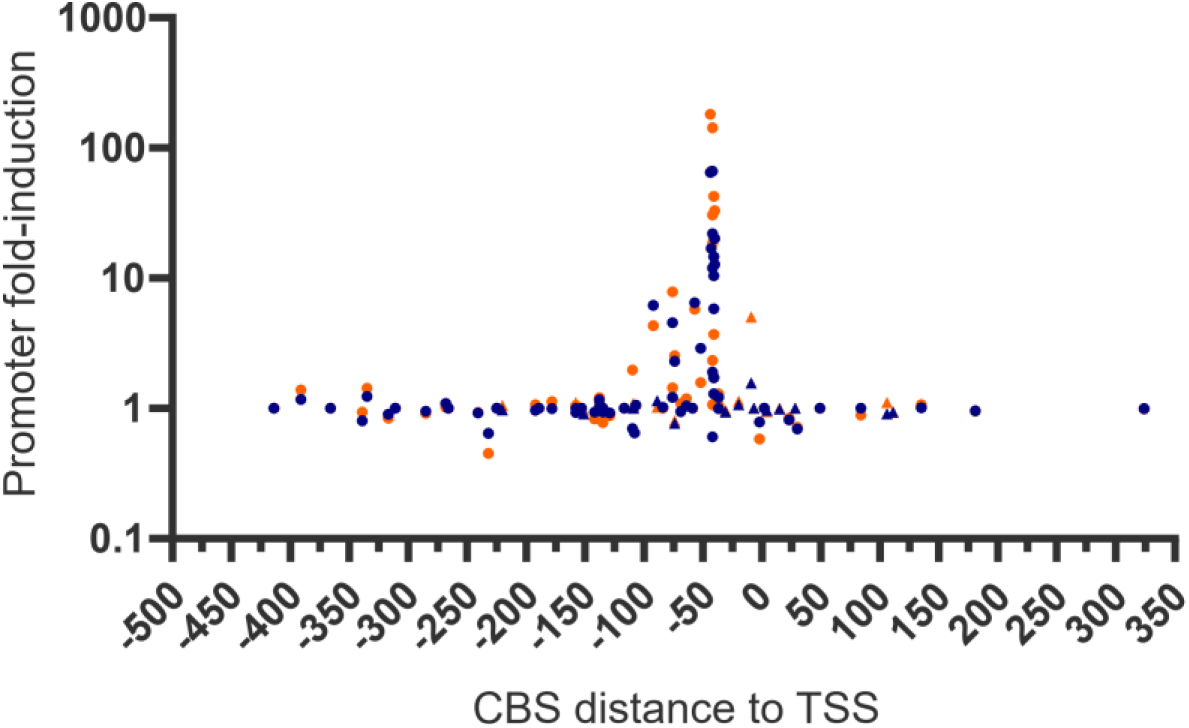
Promoter-probe induction relative to CBS location. All promoter regions with TSS data available and containing a putative CBS with a perfect (circle) or imperfect (one mismatch) (triangle) consensus sequence are displayed. TSS were derived from RNA-seq data of *S. meliloti* wild type strains (32, 33) and of the *ecf* ^−^ *S. meliloti* strain RFF625c (88). Promoter induction ratios derived from cultures supplemented with cAMP (blue) or cGMP (orange) are shown in Fig. S2E and Table S2.

### Clr affinity for cGMP increases in presence of DNA

As EMSA and promoter activation studies suggested cNMP-inducible binding of CBS-DNA by Clr, we further analysed the binding behaviour of Clr using isothermal titration calorimetry (ITC, Fig. S3). A synthetic 33 base pair DNA oligomer was designed based on the core CBS motif as binding target for Clr. It contains a double strand break and was subsequently used in crystallisation experiments (see Fig. 7). The obtained thermograms are consistent with a simple model assuming one binding site per Clr molecule, i.e., no cooperativity between the binding sites. A similar behaviour has been described for *E. coli* CAP (18, 37, 38) and *M. tuberculosis* CRP (38). The ‘active’ form of CAP is represented by a CRP dimer·cAMP2 complex (37, 39). The secondary binding site proposed for *E. coli* CAP and *M. tuberculosis* CRP located in the *C*-terminal HTH DNA binding domain is not conserved in Clr and no binding could be observed in ITC, HDX-MS and crystallographic experiments (40). As the affinity for the secondary binding site is considerably above physiological concentrations its biological relevance has recently been dismissed (9, 39, 41, 42).

The affinity of cAMP binding by Clr is mostly not affected by the presence of cognate CBS-DNA with KD values of 6-7 μM (Table 1). Formation of the ternary Clr•cAMP•CBS-DNA complex is enthalpically driven with an opposed loss of entropy, whereas that of the binary Clr•cAMP complex relies both on enthalpic and entropic contributions. In contrast, cGMP binding by Clr alone proceeds endothermically and with apparently lower affinity (KD ∼24 μM). Compared to the Clr•cAMP•CBS-DNA complex, ternary complex formation with cGMP gives similar affinities, and also relies on both contributions. For Clr alone, no DNA binding could be detected under ITC conditions. In the apo state of *E. coli* CAP, the cAMP and CBS-DNA binding domains appear as quasi-independent dynamic units, while the intra- and inter-subunit correlations increased in the holo state (43). Our ITC data indicate a profound difference only for the binary Clr•cNMP complexes, which may affect formation of ensembles capable of binding to CBS-DNA. According to size exclusion chromatography, apo-Clr forms a monomer-dimer equilibrium in solution that favours the monomeric state with 86%. For comparison, the best studied Clr homolog is *E. coli* CAP, which is able to bind both cAMP and cGMP, but is only activated by cAMP for DNA binding (44). The lack of activation in response to cGMP has also been found for *M. tuberculosis* Rv3676 and has been assumed to be a generic trait of Crp-like proteins (41).

**TABLE 1.**
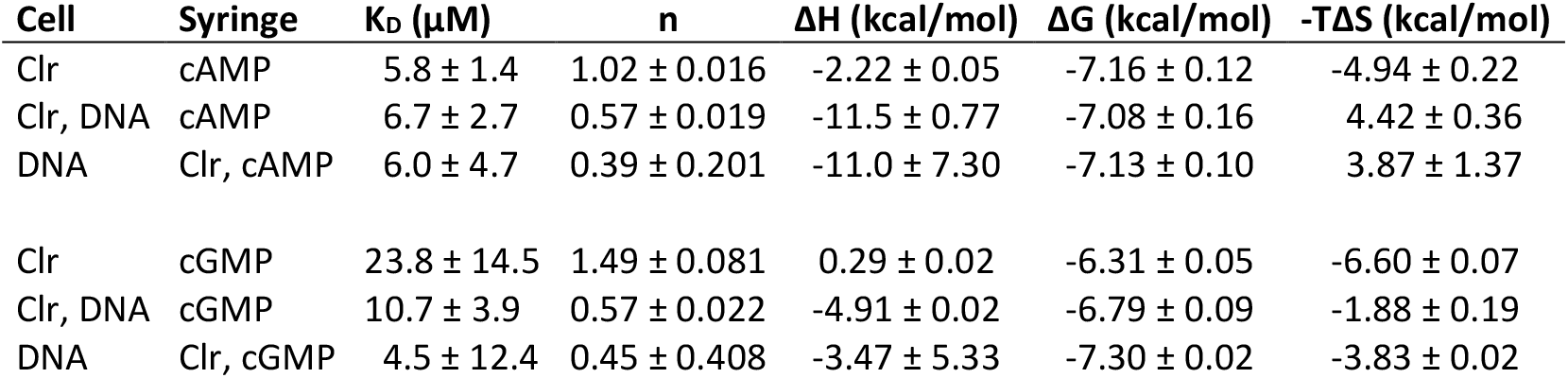
Isothermal titration calorimetry measurements for the complex assembly. The measurements were conducted at 25°C. The corresponding raw data can be found in Fig. S3.

| Cell | Syringe | $K_D$ (μM) | n | $\Delta H$ (kcal/mol) | $\Delta G$ (kcal/mol) | $-T\Delta S$ (kcal/mol) |
| --- | --- | --- | --- | --- | --- | --- |
| Clr | cAMP | $5.8 \pm 1.4$ | $1.02 \pm 0.016$ | $-2.22 \pm 0.05$ | $-7.16 \pm 0.12$ | $-4.94 \pm 0.22$ |
| Clr, DNA | cAMP | $6.7 \pm 2.7$ | $0.57 \pm 0.019$ | $-11.5 \pm 0.77$ | $-7.08 \pm 0.16$ | $4.42 \pm 0.36$ |
| DNA | Clr, cAMP | $6.0 \pm 4.7$ | $0.39 \pm 0.201$ | $-11.0 \pm 7.30$ | $-7.13 \pm 0.10$ | $3.87 \pm 1.37$ |
| Clr | cGMP | $23.8 \pm 14.5$ | $1.49 \pm 0.081$ | $0.29 \pm 0.02$ | $-6.31 \pm 0.05$ | $-6.60 \pm 0.07$ |
| Clr, DNA | cGMP | $10.7 \pm 3.9$ | $0.57 \pm 0.022$ | $-4.91 \pm 0.02$ | $-6.79 \pm 0.09$ | $-1.88 \pm 0.19$ |
| DNA | Clr, cGMP | $4.5 \pm 12.4$ | $0.45 \pm 0.408$ | $-3.47 \pm 5.33$ | $-7.30 \pm 0.02$ | $-3.83 \pm 0.02$ |

### Both cAMP and cGMP are able to shift Clr to its active conformation

Unlike apo-Clr and its binary complexes, we found that ternary Clr•cNMP•CBS-DNA complexes crystallize within a few hours in the same, orthorhombic crystal form. Given their intrinsic high anisotropic diffraction characteristics we employed extensive optimisation screening to yield crystals diffracting to 2.8 and 3.1 Å for the cAMP- and cGMP-bound Clr•CBS-DNA complex, respectively (PDB accession code 7PZA and 7PZB, Table S5). The structures show a Clr dimer that is bound to 26 bp CBS-DNA containing a central double strand break. The individual subunits fold into two domains linked by helix α5 (residues P123-L149), which is the Clr dimerization interface. The α5 helices of the dimer form a coiled-coil mainly driven by hydrophobic interactions, with T140 and T141 as notable exceptions. Residues M1-Q121 belong to the cNMP binding domain, the larger of the two. It is formed by four α helices and seven β strands in an antiparallel orientation of a β-sandwich encasing the cNMP binding site. A helix-turn-helix (HTH) DNA binding domain consisting of four α helices and two β strands is located at the C-terminus (Y150-D234). The cNMP binding domain is universally conserved throughout cNMP receptor proteins, as well as protein kinases and ion channels, with sequence identities above 20% even for mammal representatives (17). The HTH motif of the DNA binding domain is prevalent in 116 structures so far deposited in the PDB according to InterPro classifications (45, 46). Thus, both domains perform their function as core components of a variety of proteins adapted to fulfil different biological tasks. Accordingly, the overall fold of Clr is highly similar to its *E. coli* homologue, the catabolite activator protein CAP (1CGP) (47) and even more so to the *M. tuberculosis* (3MZH) and *C. glutamicum* CRPs (4CYD) (48, 49). The latter two served as templates for molecular replacement.

The CBS-DNA interacts with its symmetry mate in the crystal lattice and is therefore well resolved. When comparing the ternary complexes, the Clr•cNMP structures as well as the CBS-DNA show a high conformational congruity with pairwise r.m.s.d. values below 0.6 Å. This further supports our notion that both cyclic nucleotides shift ternary Clr•cNMP•CBS-DNA complexes to almost identical active conformations. The cNMP binding site located in the N-terminal domain is stoichiometrically occupied by cAMP and cGMP, respectively (Fig. 6A and B, and Fig. S4). The binding mode of the phosphate and sugar moieties is identical, involving G85 and E86 within α4 as well as R95 and S96 of the following loop. The sidechain of S96 is flipped towards the phosphate only in the cGMP-bound complex. The nucleobase of cGMP is flipped to the *syn-*conformation, while it occupies the *anti*-conformation in the Clr•cAMP complex. Both nucleobase conformations allow for interaction with helix α5 residue T140 of the same Clr subunit and T141 of the adjacent Clr subunit. While cAMP interacts via its primary N^6^-amine with both T140 and T141, cGMP forms hydrogen bonds between T140 and its C6 carbonyl group as well as between T141 and the N^2^-amine. The latter interaction between the N^2^-amine of cGMP and T141 of the adjacent molecule is a substantial difference between Clr and the inactive cGMP complex of *E. coli* CAP (50). Interaction of the cAMP nucleotide with both α5 helices has been described to trigger a coil-to-helix transition in the *C-*terminal part of the hinge and shift the HTH domain towards its active conformation for CAP (51). So far, cGMP appeared to be incapable of such an interaction with the second helix for promoting CRP activation (44). In case of Clr both the interaction and arrangement of the full α5 helix is resolved by the structure, which is in an agreement with Clr-DNA binding *in vitro* and activation of promoter activity *in vivo* by both cNMPs. Moreover, mapping the relative hydrogen-deuterium exchange between apo and holo states by HDX-MS experiments shows that upon nucleotide addition the *C*-terminal half of the helix gets shielded from exchange. This further supports the notion, that both nucleotides are capable to trigger the coil-to-helix transition in the linker to the C-terminal helix-turn-helix domain (Fig. 6C and D). This shielding is not only due to nucleotide interactions with T140 and T141, as even fragments, containing the last five residues of the α5 helix, show a clear decline in solvent accessibility for nucleotide-bound states. Helix α6 is shortened in response to the hinge rearrangement and interacts with the former in an angle of about 40° in the active conformation (11). This interaction is stabilised in Clr by a formerly unknown chloride-binding site. The relative hydrogen-deuterium exchange comparing the apo and holo states also shows a slightly stronger decline in deuterium uptake within the binding pocket for the cAMP-bound compared to the cGMP-bound state. Some uptake differences within helix α7 and α8 of the HTH domain can be observed (Fig. S4). Upon nucleotide binding the deuterium exchange increases in this region, indicating some degree of conformational change. This is in agreement with structures published for *E. coli* CAP, where the active conformation is achieved by a 60° rotation of α8 (44, 52, 53).

**FIG 6.**
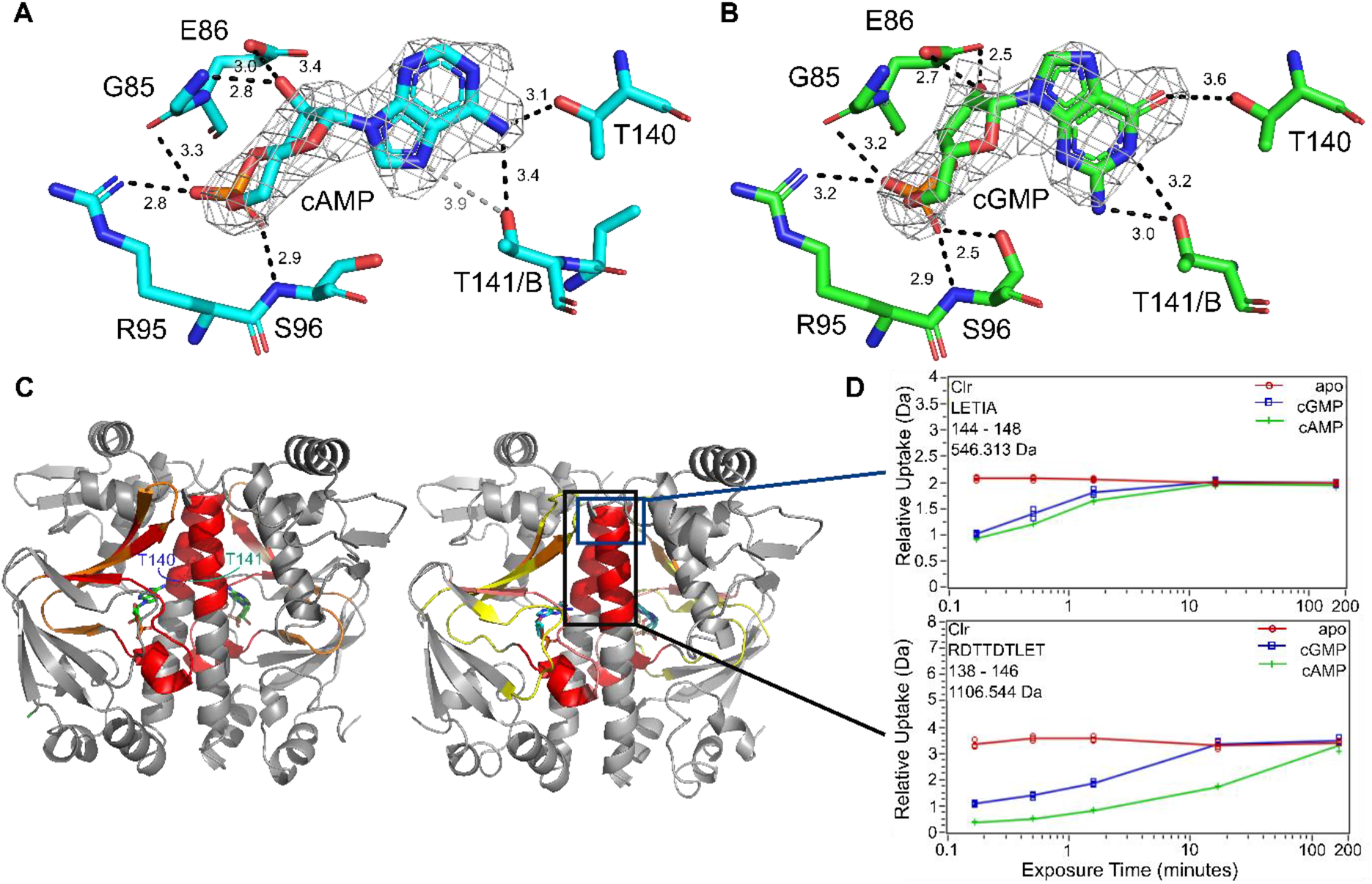
Characteristics of the Clr-ligand interaction. (A, B) Nucleotide binding environment in the cAMP- and cGMP-bound crystal structures. The sugar and phosphate moiety of the ligand interact with G85 and E86 of helix α4, as well as R95 and S96. The sidechain of Ser96 is tilted towards the ligand in the cAMP-bound structure and Thr141 of the adjacent subunit is in closer proximity to it.; The primary amine of the adenosine is coordinating Thr140 from α5 in the same subunit and Thr141 from the adjacent one. The ligand 2 m*F*o−D*F*c experimental electron density maps contoured at 1.3 σ are shown in grey. Binding interactions are indicated in Å. (C) Areas of reduced relative hydrogen-deuterium exchange upon binding of cAMP (left) and cGMP (right) coloured in from 5-20% in 5% increments (yellow, orange, red). In addition to the expected displacement of solvent in the binding cavity, the *C*-terminal half of helix 5 (black box) shows a reduction of exchange, that can be attributed to a coil-to-helix transformation. (D) Time-dependent deuterium exchange graphs for two key peptides within helix 5 show, that the shielding cannot be solely due to interaction with T140 and T141, but is the result of a change in secondary structure. Peptides not including both exemplified by ^144^LETIA (blue box) also show a decrease in H/D exchange.

### The crystal structures of nucleotide bound Clr shed light on its DNA specificity

Clr binds to a target DNA of 33 bp length via electrostatic interactions and hydrogen-bonding to the major groove. The surface of the HTH DNA binding domain has a strongly positively charged patch with which it interacts with the DNA phosphate backbone (Fig. 7A). Clr has a likewise positively charged cavity in each subunit for accommodating cNMP. Helix α8 is inserted in the major groove and interacts with hydrogen bonds over six bases as depicted in Fig. 7B and C. The binding results in a 40° and 32° kinking of the DNA helix at the longer and shorter DNA oligomers, respectively, which is eased by the engineered double strand break at the twofold symmetry centre. The entropic cost associated with DNA bending is overcompensated by establishing further interactions. As a result, the flexibility of the flanking regions of the recognition sequence might be distinguished by Clr, adding another layer of specificity (54, 55).

The electron density of the duplex DNA that was annealed from four oligonucleotides (Fig. 7D) defines the 14 bp of the 5’-oligonucleotides of the core as well as 19 (17) bp of the two 3’-oligonucleotides. The base readout occurs at most conserved positions of the core CBS motif (Fig. 7D) (23). Q184 of α7 and R195 and N199 of α8 are interacting with the ^9^TGT triplet of the 14-mers, whereas K197 and S194 of the same helix, as well as L152 within the loop *C*-terminally of α5 are hydrogen-bridged to the ^4^GGT triplet of the 19-mer. For *E. coli* CAP mainly R180, E181 and R185 have been identified to interact with the recognition sequence (56, 57). In Clr, R180 corresponds to R195, E181 is not conserved and R185 is conserved as R200, but fails to interact with the DNA as it adopts a different rotamer. Accordingly, CRP-like proteins in the same phylogenetic subclass as *E. coli* CAP have a highly conserved RE(T/M)VGR motif in their α8 helix, whereas CRP subclass G, to which Clr belongs, is significantly more diverse in this region (16).

**FIG 7.**
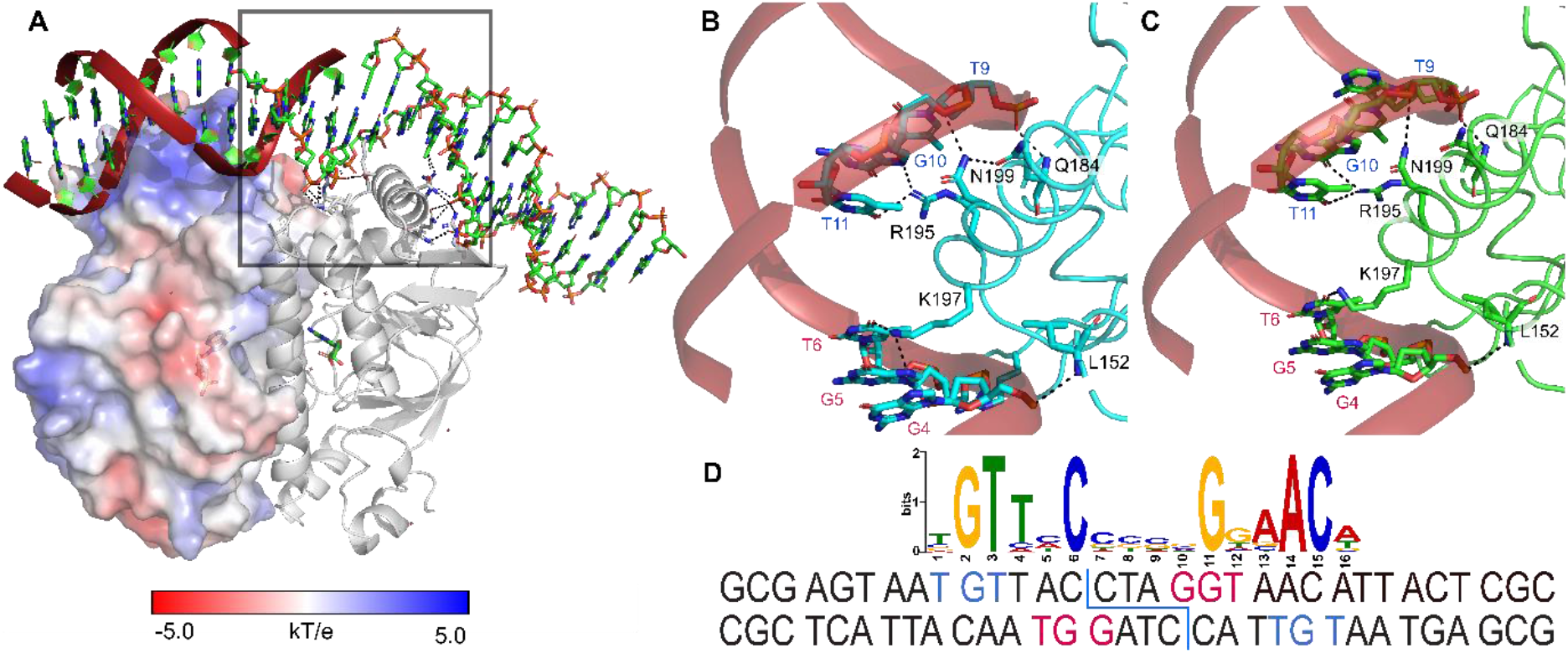
Interaction of Clr with its DNA recognition site. (A) The surface of the helix-turn-helix DNA binding domain is positively charged to interact with the phosphate backbone. Helix α8 is inserted into the major groove and provides base readout via direct hydrogen bonding interactions. The double strand break above the subunit interface facilitates the DNA kinking of about 40°. (B, C) Direct interactions of Clr with the DNA major groove upon cAMP (B) and cGMP (C) binding. The DNA bases involved in the interaction are highly conserved. (D) The synthetic oligonucleotide used in crystallisation has a double strand break with sticky ends right above the subunit interface; the 5’-oligonucleotides of the annealed core motif (GCGA…) belong to chains D and F, the 3’-oligonucleotides to chains C and E of the structure. The design was based on the consensus motif GTNNCNNNNGNNAC.

CRP proteins interact with the bacterial RNAP via the stacked β-strands within the HTH domain that has been termed AR1 in class I promoters (56, 58) or the top of the beta barrel within the cNMP binding domain, which has been labelled AR2, in class II promoters (59, 60). For Clr AR1 three of the four beta strands are present, whereas the N-terminal loop of the motif is turned more towards the protein core (Fig. S4). Crystal structures of CAP in complex with DNA and the αCTD domain of the RNAP show very little flexibility within the interface and surprisingly only one hydrogen bond between T158 of CAP and T285 of the αCTD (PDB accessions 5CIZ, 3N4M and 1LB2). The interface is largely hydrophobic, a feature that is retained in Clr. The activating region AR1 has a significantly divergent orientation in its *N*-terminal part for Clr. This does prevent interaction with the *E. coli* RNAP *in vivo*, which we infer from promoter activation by Clr in *E. coli* (see above). The overall conservation of the 22 residues comprising AR1 is rather low with three identical and three similar amino acids. In class II promoters the RNAP interacts with the CRP on two additional sites: AR1 of the second dimer symmetry unit and AR2 of this unit (3). The interactions with AR2 involve the beta and omega subunit as shown in crystal and cryo-EM structures from *Thermus thermophilus* and *E. coli* (60, 61). The interactions of Clr within the omega unit are conserved as G47 and D48 (G14 and E15 in *T. thermophilus*), whereas the area involved in the interaction with the beta subunit in *T. thermophilus* (mainly through E8) deviates in a way that allows no deductions on whether it would interact in the same way.

## DISCUSSION

Our study was focused on the bifunctionality of Clr. We confirmed that Clr is able to bind cAMP and cGMP *in vitro* and characterized the binding properties of this transcription regulator for both these cyclic nucleotides and for its DNA binding site. In our ITC measurements cAMP bound to Clr with about four times higher affinity than cGMP (K_D_^cGMP^ = 24μM; K_D_^cGMP^ = 6 μM). For CAP from *E. coli*, values in the same range have been reported, the affinity for cAMP (10 μM) also being greater than that of cGMP with 16 μM (9, 40). However, we showed that the difference between the nucleotide affinities of Clr diminished as soon as cognate DNA was present, showing that ternary complex formation by both effector molecules is similarly driven.

The regulation of cAMP receptor proteins depends on an equilibrium between an active form that is capable of DNA interaction and an inactive form that is not. The equilibrium is shifted towards the active conformation by effector binding (18). For Clr, both our crystal structures and the HDX experiments clearly prove that Clr undergoes this switch independently of the purine base of cNMP. As the cNMP binding site does not deviate significantly from that of the *E. coli* homologue, Clr might have a lower energy barrier for the shift in population towards the active conformation. This is further supported as similar equilibrium shifts have been introduced via mutations of *E. coli* CAP before (62). Cyclic AMP binding initiates the population shift by interacting with residues of the central helices of both subunits (CAP:T127 and CAP:S128) (63). This in turn leads to a coil-to-helix transition in the *C*-terminal part of the helix, resulting in a DNA binding domain in its active conformation (11, 52, 64, 65). Contrary to any previous findings in *E*. coli, we clearly established that Clr performs an identical binding movement via a ^140^TT motif for both cNMPs. As cGMP is bound in a *syn* conformation the interactions can be retained. We were able to show the resulting coil-to-helix switch of the α5 helix by HDX-MS in solution due to a decline in deuterium uptake, further proving the bifunctional nature of Clr. Clr residues G85, E86, R95 and S96, which are also part of the binding pocket, are largely conserved in CAP as well and responsible for the high-affinity nucleotide binding (66).

The C-terminal helix-turn-helix motif of CRPs recognizes its cognate consensus DNA motif via major groove readout (43). Binding of the Clr dimer to two symmetrically related motifs results in a kink of the DNA by 40° and 32°. For CAP, DNA kinks of even 110° have been reported and modified promoters containing an inherent kink were shown to function CRP-independent, which highlights the importance of this particular feature (67, 47, 57, 68, 69). Our isothermal titration calorimetry measurements of DNA binding gave dissociation constants of 6.0 and 4.5 μM and free energy changes of -7.1 and -7.3 kcal/mol for cAMP and cGMP, respectively. This makes Clr a transcription factor on the weaker side of the binding spectrum. However, stronger does not always equal better, as the binding needs to be reversible as well (54). For duplex-DNA of similar length CAP dimer·cAMP2 binding affinities of about 10 μM have been reported, which suggests similar binding strengths for CAP and Clr (69).

Clr interacts with the nucleobases of the DNA core motif via hydrogen bonds. The residues involved in sequence readout are exactly contacting the bases within the CBSs as derived from genome-wide ChIP-seq and EMSA data of this work. The bifunctionality of Clr was demonstrated *in vivo* as well. Virtually all promoters directly targeted by Clr were activated by Clr-cAMP and Clr-cGMP in both *S. meliloti* and *E. coli*. The arrangement of three overlapping imperfect CBS core motifs in the *SMc05008* upstream region is intriguing. It suggests a binding mode different from binding of a single CBS or that differential binding of the individual sites might generate a different regulatory output. In *S. meliloti*, we found that externally added cGMP compared to cAMP activated gene expression up to four times higher. Both, the affinities as well as the structures of cNMP-bound Clr showed no clear distinction explaining the higher degree of cGMP-dependent promoter activation in *S. meliloti*. Moreover, stronger cGMP-mediated activation was not observed in *E. coli*. The difference is therefore likely rooted in higher cGMP levels within the *S. meliloti* cell under the *in vivo* assay conditions used in this study. Whether this is caused by endogenous AC/GCs or differences in uptake of exogenous cNMP remains to be elucidated. Previously, it was shown that *S. meliloti* is able to produce cGMP upon overproduction of its endogenous AC/GC CyaB (23).

However, which cNMP molecules are synthesized under native conditions is not yet known, as concentrations seem to be rather low. Except *cyaF2*, which encodes a putative AC or GC, all other identified genes directly regulated by Clr-cNMP code for hypothetical proteins. Eight of the Clr-cNMP regulated genes encode small proteins of up to 100 amino acids, six of these with an *N*-terminal signal peptide. *SMb20495, SMc02177* and *SMc02178* predicted to encode hypothetical proteins were previously shown to participate in autoregulation of alfalfa root hair infections (5, 30, 70). With exception of these three genes, the biological role of the other identified Clr-cNMP targets remains unclear. Because many of the direct Clr-cNMP controlled target genes encode proteins with *N*-terminal signal peptide, we speculate that most of these proteins might be relevant for extracellular functions and possibly involved in the symbiotic interaction with the host plant.

Notably, Clr-cNMP was found to directly induce transcription of the small non-coding RNA *SMelC181* of unknown function. Only a few studies reported regulation of small non-coding RNAs by a CRP. Examples are small non-coding RNAs associated to the regulation of quorum sensing, nitrogen assimilation, and galactose utilization in *E. coli, Salmonella, Vibrionales* and cyanobacteria (71-73). This makes *SmelC181* an interesting candidate for upcoming studies.

In the last two decades, cNMP signaling networks have been characterized in several bacterial species and different CRPs have been studied for their cNMP and DNA binding properties (3, 6, 10, 74). These studies imply that cNMP signaling networks and CRPs have evolved differently in the context of the different habitats of these bacteria. In this study, we showcased a CRP-like protein that has evolved to be activated both by cAMP and cGMP to perform its function in transcriptional regulation in a plant-symbiotic α-proteobacterium. The biological function of Clr bifunctionality for binding of these two cNMPs remains to be discovered. For example, it might have evolved as an adaptation to endogenous bacterial cGMP synthesis or for sensing of plant-derived cGMP. Although the *S. meliloti* Clr regulon has been extensively characterized, current knowledge of its biological functions is limited to its role in early stages of the symbiosis (5). Deciphering the biological function of the responsiveness of Clr to cAMP and cGMP and whether subclass G CRP transcription factors are generally characterized by being activatable by cGMP or by both cGMP and cAMP is an exciting goal of future research.

## MATERIALS AND METHODS

### Strains, plasmids and growth conditions

Unless specified otherwise, *E. coli* was grown at 37°C in LB-Lennox medium (1% tryptone, 0.5% yeast extract, 0.5% NaCl) with the addition of kanamycin (50 μg/mL), gentamycin (8 μg/mL or tetracycline (10 μg/mL) as needed. *S. meliloti* was grown in TY (0.5% tryptone, 0.3% yeast extract, 2.7 mM CaCl2) or MOPS minimal medium (1% MOPS, 1 mM MgSO4, 20 mM sodium glutamate, 20 mM mannitol, 2 mM K2HPO4, 250 μM CaCl2, 37 μM FeCl3, 4.1 μM biotin, 45.8 μM H3BO3, 10 μM MnSO4, 1 μM ZnSO4, 0.5 μM CuSO4, 0.27 μM CoCl2, 0.5 μM Na2MoO4). If required, streptomycin (600 μg/mL), kanamycin (200 μg/mL), gentamycin (30 μg/mL) or tetracycline (10 μg/mL) were added to solid medium containing agar as solidifying agent. For liquid media, the concentration of kanamycin, gentamycin or tetracycline were reduced by half.

Plasmids were transferred to *S. meliloti* by *E. coli* S17-1-mediated conjugation as previously described (75).

Promoter-probe constructs were generated by insertion of a *S. meliloti* gene upstream region of 300 to 735 bp and up to 75 bp of the associated protein or RNA coding region into the replicative low copy plasmid pPHU231-EGFP or replicative medium-copy plasmid pSRKKm-EGFP for studies in *S. meliloti* strains. For the promoters of small non-coding RNA genes, a Shine-Dalgarno sequence was included downstream of the selected promoter sequence.

Site-directed mutagenesis of putative CBSs was performed by overlap extension PCR and mutated promoter regions of the same size as the corresponding wild type promoter region were inserted into pSRKKm-EGFP.

For cloning of CBSs, two complementary oligonucleotides (1 μM each) were hybridized in T4 DNA-ligase-buffer (NEB) containing T4 polynucleotide kinase (PNK; 0.2 μl; NEB) in a total volume of 10 μl to obtain double-stranded DNA flanked by *Xba*I and *Hind*III overhangs. Phosphorylation of the oligonucleotides was performed at 37°C for 45 min followed by a PNK deactivating step at 65°C for 20 min. The reaction was then heated at 95°C for 5 min and slowly cooled down to 30°C. 30 ng of the hybridized product was directly used as insert for ligation with pSRKKm-EGFP.

For construction of the C-terminal 3xFLAG-tagged Clr overexpression construct, the full-length *clr* coding sequence was amplified from genomic DNA of *S. meliloti* Rm2011, digested with *Nde*I and *Xba*I and inserted into plasmid pSRKKm-CF, yielding pSRKKm-*clr*-CF. Subsequently, *clr*-CF was amplified from this plasmid, digested with *Xba*I and *Hind*III and inserted into plasmid pWBT, yielding pWBT-*clr*-CF. Expression plasmids pWBT-AC/GC and pWBT-AC/GC-c*lr* are described in (23).

To obtain *S. meliloti* gene deletion mutants, gene-flanking regions of 350 to 750 bp were cloned into suicide vector pK18mobSacB. Following conjugation-mediated plasmid transfer to *S. meliloti* Rm2011 and plasmid integration into the genomic DNA by homologous recombination, transconjugants were subjected to sucrose selection as previously described (76). Gene deletions were verified by PCR. The *S. meliloti* multiple deletion mutant *cya*^*0*^ lacking all 28 putative class III AC/GC genes was generated by sequential gene deletions. The genomes of *cya*^*0*^ strains as well as the parenting strain Rm2011 were sequenced. Total DNA was purified using a DNeasy blood and tissue kit (Qiagen). DNA sequencing libraries were generated by applying the Nextera XT DNA Library Preparation kit (Illumina). Sequencing was performed on a MiSeq Desktop Sequencer (Illumina) using the MiSeq reagent kit v2, for 2 × 250 bp paired-end reads (Illumina) for Rm2011 resulting in 4.45 × 10^6^ reads or the MiSeq reagent kit v3 for 2 × 75 bp paired end reads for Rm2011 *cya*^*0*^ resulting in 5.82 × 10^6^ reads. Single-nucleotide polymorphism (SNP) detection was performed by applying CLC Genomics workbench (v10.1.1, Qiagen). Paired reads were mapped to the annotated reference genome of *S. meliloti* Rm1021 (chromosome, AL591688; pSymA, NC_003037; pSymB, AL591985). The minimum variation frequency was 50 % and at least ten reads with a minimum mismatch coverage of eight were considered. The *S. meliloti cya*^*0*^ strain used in this study (Fig. 4, Table S4) carries one SNP (Table S1B).

To construct the expression plasmid coding for C-terminally His6-tagged Clr, the native encoding sequence was amplified by PCR and inserted into *Nde*I- and *Xho*I-digested pET36b(+) (Novagen). The fusion construct was verified by DNA sequencing. Strains, plasmids, and oligonucleotides used in this study are described in Tables S6.

### EGFP fluorescence measurements

*S. meliloti* strains were grown in 100 μl TY-medium and *E. coli* BTH101 in 100 μl LB-Medium in 96-well polystyrene flat bottom plates (Greiner) at 30°C shaking at 1200 rpm. For cultures in which cNMPs were added to the medium, stationary pre-cultures of the respective strains grown in TY-medium without cNMPs were diluted 1:100 in medium containing 400 μM cAMP or cGMP, or as control in medium without cNMPs, followed by incubation for 24 h. For promoter studies in *E. coli* BTH101, stationary pre-cultures were diluted 1:100 in medium containing 100 μM IPTG followed by incubation for 16 h.

EGFP fluorescence measurements were carried out as described in (23). Relative fluorescent units (RFU) represent fluorescent values divided by the optical density (OD600). The measurements were carried out in *S. meliloti* Rm2011, Rm2011 Δ*clr*, Rm2011 *cya*^*0*^ and *E. coli* BTH101 background. Processed data and underlying raw data of each experiment are listed in Tables S2-13.

In *S. meliloti* strains, the background fluorescence of the corresponding empty vector control was subtracted (Fig. 1 A and C, Fig. S2E, Tables S2 and S4). In measurements carried out in *E. coli* BTH101 (Fig. 3 B, Table S3), no background fluorescence was subtracted. Fluorescence fold-change of *S. meliloti* strains carrying promoter-probe constructs was calculated from the RFU of the induced strains divided by the RFU of the non-induced strains or by the RFU of the wild type divided by the RFU of the mutants (Table S2). In *E. coli*, fluorescence fold-change was calculated by the fluorescent values derived from the strains carrying pPHU231-promoter-probe-EGFP in combination with pWBT-AC/GC-*clr*-EGFP divided by the values derived from the strains carrying pPHU231-promoter-probe-EGFP in combination with pWBT-AC/GC-EGFP (Table S3). Three to four independent transconjugants and transformants of each strain containing the promoter-*egfp* constructs were used as biological replicates.

### ChIP-Seq

Cultures of Rm2011 Δ*clr* carrying the *clr*-FLAG overexpression construct pWBT clr-CF were grown in 60 ml of TY or MOPS in 500 ml flasks each supplemented with either 400 μM 3’,5’-cAMP or 3’,5’-cGMP and 500 μM IPTG. Upon reaching an OD600 of 0.6, cells were fixated with 1% formaldehyde for 20 minutes at room temperature (RT). Fixation was quenched with 250 μM glycine for 20 minutes.

Cell-pellets were resuspended in 300 μl IP-buffer (50 mM HEPES-KOH pH7.8; 150 mM NaCl, 1 mM EDTA, 1% Triton X100, 0.1% sodium deoxycholate, 0.1% SDS) with 1 mM of phenylmethylsulfonyl fluoride (PMSF). Cells were lysed and DNA fragmented (∼500 bp) by sonication in a Bioruptor® Plus (Diagenode) for 48 × 30 s and 30 s cooling. 300 μl sonicated product was mixed with 3 ml IP-buffer and 1 mM PMSF and centrifuged at full speed at 4°C for 30 min. 100 μl supernatant were taken and frozen at -20°C as a control. Clr-CF was immunoprecipitated with ANTI-FLAG® M2 Affinity Gel (Sigma-Aldrich). The protein was eluted with 100 μl 3x FLAG peptide elution buffer for 1 h at 4°C. Cross-links of control and ChIP-DNA were removed with 200 μM NaCl and proteins and RNA were degraded with proteinase K and RNAse A overnight at 65°C. Samples were checked in a Western blot with monoclonal anti-FLAG M2-peroxidase (HRP) antibody (1: 1000). DNA was purified with the QIAquick PCR kit. 10 ng DNA was taken for ChIP-Seq DNA library preparation and 2.5 ng for qPCR reaction. The DNA library preparation was carried out as described in (77) with the adapter sequences 2, 4-7 and 12-14 compatible with Illumina’s TruSeq platform. Sequencing was performed on a MiSeq Desktop Sequencer (Illumina) using a MiSeq reagent kit v3 with 2 × 75 paired end-reads and sequence analysis was performed using CLC-Genomics workbench (v10.1.1., Qiagen). Enrichment in reads of each sample was compared to the corresponding control and mapped as peaks against the annotated genome of *S. meliloti* Rm1021. Sequences flanking each center of peak derived from the ChIP-Seq analysis were defined and summarized as FASTA sequences using the R package (78). Motifs were generated with the MEME Suite online tools MEME (v5.1) and FIMO (v5.3) (79, 29).

### Electrophoretic mobility shift assay

An electrophoretic mobility shift assay (EMSA) reaction mixture contained 2 mM HEPES (pH 7.0), 30 mM NaCl, 50 mM KCl, 850 ng sonicated salmon sperm DNA (GE Healthcare), 1 μg of BSA (Sigma) and 20 ng of Cy3-labeled DNA in a final volume of 10 μl. Native and synthetic Cy3-labeled DNA fragments were obtained by PCR with Cy3-labeled primers (Table S15D) using the corresponding pSRKKm-promoter-EGFP or pSRKKm-CBS-EGFP constructs as template, respectively. 592 bp synthetic Cy3-labeled DNA fragments were composed of the CBS flanked by pSRKKm-EGFP-derived sequence. The protein was added at 3 μg (111 μM) per reaction and 3’,5’-cAMP or 3’,5’-cGMP at 1 mM if indicated. The reaction mixtures were incubated at room temperature for 30 min in the dark. A mixture of 1 μl of 90% glycerol and 1.5 μl of 5x TBE buffer was added to each reaction, and 10 μl of which was loaded onto a 10% polyacrylamide gel in 1x TBE. Following electrophoresis at 9 V cm^−1^ at room temperature for 3.5 h, images were taken using a Typhoon 8600 variable mode imager (Amersham Bioscience). Sequences of every DNA fragment used in this study are listed in Table S15B and C.

### Protein expression and purification

Clr was overproduced in *E. coli* BL21 (DE3) cells in LB-Lennox medium containing kanamycin (30 μg/mL) at 37°C and induced at an optical density of 0.5 (600 nm) using IPTG (0.1 mM). After 3 h after addition of 0.1 mM IPTG and harvested by centrifugation (15 min at 4°C; 4000 rpm). The pellet was resuspended in 20 mL binding buffer (50 mM sodium dihydrogen phosphate, 1000 mM sodium chloride, 10 mM imidazole, pH 7.0) and disrupted by three passages through a cold French pressure cell press at 15.2 bar cm^−2^. The lysate was centrifuged at 18000 rpm for 60 min at 4°C and filtered through 0.45 μm membrane filter. For Clr purification the initial purification was conducted using a 5 mL Protino™ Ni-NTA Column (Macherey-Nagel). Size-exclusion chromatography over a 16/600 Superdex 200 column (GE) using elution buffer (20 mM HEPES, 300 mM NaCl, pH 7.0) was done as a polishing step. Preceding the purification, the size-exclusion buffer was optimized as described using SYPRO™ Orange dye (89).

### Protein crystallization

Clr (0.25 mM/0.44 mM) was crystallized in the presence of 25 mM cAMP/cGMP, 25 mM MgCl2 and a 1.25 times excess of 19-mer/14-mer duplex target DNA (19-mer sense strand: 5’-CTA GGT AAC ATT ACT CGC-3’; 14-mer antisense strand: 5’-GCG AGT AAT GTT AC-3’). The oligonucleotide (BioCat) was prepared from single strands by heating equimolar amounts to 98°C for 5 min and slowly cooling to room temperature. The DNA oligonucleotides used for the crystallisation of Clr were designed based of the genes Smc04190 and SMc00925, which have previously been reported to be regulated by Clr (23).

Crystals of Clr in complex with cAMP were grown at 291 K by sitting-drop vapour diffusion in 0.09 M sodium fluoride, 0.09 M sodium bromide, 0.09 M sodium iodide, 0.1 M Tris base/BICINE pH 8.5, 12.5% (v/v) MPD, 12.5% (w/v) PEG1000 and 12.5% (w/v) PEG3350. Crystals containing cGMP were grown at 281 K in sitting-drop vapour diffusion plates in 0.2 M 1,6-hexanediol, 0.2 M 1-butanol, 0.2 M (*RS*)-1,2-butanediol, 0.2 M 2-propanol, 0.2 M 1,4-butanediol, 0.2 M 1,3-propanediol, 0.1 M MOPS/HEPES-Na pH 7.5, 12.5% (v/v) MPD, 12.5% (w/v) PEG1000 and 12.5% (w/v) PEG3350.

The crystals were flash-frozen in liquid nitrogen and diffraction data collected at 100 K at the EMBL/DESY P13 beamline (Hamburg) using a Pilatus 6M detector (Dectris) and SLS PXI X06SA beamline (Paul-Scherrer Institute, Villigen) using an Eiger 16M detector (Dectris), respectively. At wavelengths of 0.976/1.0 Å the crystals diffracted to 2.8 and 3.1 Å. The data were processed using XDS (81) in Space Group P21 21 21. Data reduction and scaling were done using CCP4i2 (v7.0.065) (82), AIMLESS in particular (Version 0.7.3). The phases were solved by molecular replacement using a hybrid model derived from GlxR from *Corynebacterium glutamicum* (PDB 4CYD) and Crp from *Mycobacterium tuberculosis* (PDB 3MZH). The model was built using COOT (v0.8.9) (83) and refinement in Phenix (v1.11.1) (84). Final refinement statistics are given in Table S5. The coordinates and structure factors were deposited to the Protein Data Bank under PDB codes 7PZA and 7PZB.

### ITC measurements

Clr was transferred to the binding buffer (20 mM HEPES, 300 mM NaCl, 20 mM MgCl2, pH 7.0) via a PD-10 column (Cytiva Life Sciences) and the titration calorimetry measurements were performed at a Malvern MicroCAL PEAQ-ITC as previously described by (85). A typical titration consisted of injecting 2 μl aliquots of 5 mM of the ligand solution into 0.3 to 0.4 mM of the protein solution every 2.5 min to ensure that the titration peak returned to the baseline prior to the next injection. For the measurement of DNA affinities, the setup was reversed, titrating protein at 0.3 to 0.4 M containing 1 mM of cyclic nucleotide into DNA at 25 μM. The cell was temperature-controlled to 25°C. Titration curves for the dilution of the ligand solution were deducted from the data.

### HDX-MS measurements

Sample preparation is automated with a two-arm robotic autosampler (LEAP Technologies). 7.5 μL of 50 μM Clr with and without 1 mM cNMP were mixed with 67.5 μL D_2_O-based SEC buffer. After 10, 30, 95, 1000 and 10000 s at 25°C respectively the H/D exchange was quenched by addition of equal parts quench buffer (400 mM KH2PO4/H3PO4, 2 M guanidine hydrochloride, pH 2.2) at 1°C. The solution was injected into an ACQUITY UPLC M-class system with HDX technology (Waters). The protein was digested online with immobilized porcine pepsin at 12°C at 100 μL/min (H2O, 0.1% formic acid), and the resulting peptides were collected on a trap column (2 mm × 2 cm) with POROS 20 R2 material (Thermo Scientific) at 0.5°C. After 3 min, the trap column was switched online with an ACQUITY UPLC BEH C18 1.7 μm 1.0 × 100 mm column (Waters), and the peptides were eluted at 0.5°C using a gradient of (A) H2O, 0.1% formic acid and 1 acetonitrile, 0.1% formic acid at 30 μL/min (5 to 35% B in 7 min, 35% to 85% B within 1 min and isocratic flow 85% B for 2 min). The column was washed for 1 min at 95% B and equilibrated at 5% B for 5 min after this. Peptides were ionized by electrospray ionization at 250°C source capillary temperature and a spray voltage of 3.0 kV. Mass spectra were acquired on a G2-Si HDMS mass spectrometer with ion mobility separation (Waters) over a range of 50 to 2,000 m/z in HDMSE or HDMS mode for undeuterated and deuterated samples, respectively. Lock mass correction was performed with [Glu1]-fibrinopeptide B standard (Waters). Between samples, the pepsin column was washed three times with 80 μL of 4% (vol/vol) acetonitrile and 0.5 M guanidine hydrochloride, and additionally, blank runs were performed between samples. Peptides were identified, and deuterium uptake was determined employing the PLGS and DynamX 3.0 software suites (both Waters) as described by (86).

## Data availability

The ChIP-seq data have been deposited in the ArrayExpress database at EMBL-EBI (www.ebi.ac.uk/arrayexpress) under the accession number E-MTAB-11788 (87). TSS data of S. meliloti strain RFF625c are available from ArrayExpress (E-MTAB-12127). The Clr protein crystal structure data has been deposited to the Protein Data Bank archive (PDB) under accession codes 7PZA and 7PZB. All study data are included in the article and/or SI Appendix.

## ACKNOWLEDGMENTS

The authors thank Ralf Poeschke for technical support during crystallization (MarXtal facility, University of Marburg), Torsten Waldminghaus for providing the ChIP-Seq DNA-library adapter and the Bioruptor Plus (Diagenode), Lotte Søgaard-Andersen for providing the Typhoon 8600 variable mode imager (Amersham Bioscience), Bernadette Boomers for technical support of ChIP-seq (Screening and Automation Technology facility, SYNMIKRO Technology Platform, University of Marburg), Patrick Sobetzko for providing the DNA-Sequence generating R-script, and staff of the beamlines P13 as operated by EMBL Hamburg at the PETRA III storage ring (DESY, Hamburg, Germany) and X06SA of the Swiss Light Source (SLS, Villigen, Switzerland). This work was supported by grants of the German Research Foundation (DFG) to LOE (ES152/14) and AB (BE2121/8) and by the DFG core facility for interactions, dynamics and macromolecular assembly structure. We thank Wieland Steinchen for HDX-MS data acquisition and assistance with HPLC measurements.

## SUPPLEMENTAL MATERIAL

**FIG S1** Functionality of Clr-CF.

**FIG S2** EMSA shift patterns of all putative Clr binding sites and promoter-probe measurements in *S. meliloti*.

**FIG S3** Raw and integrated ITC thermograms.

**FIG S4** Structural features of DNA binding by Clr.

**TABLE S1** ChIP-seq data and analyses.

**TABLE S2** Clr-dependent promoter-probe activation.

**TABLE S3** Clr-cNMP dependent promoter-probe activation *E. coli* BTH101.

**TABLE S4** EGFP-derived fluorescence fold-reduction in *S. meliloti* Δclr and cya0.

**TABLE S5** Data collection and refinement statistics for cNMP-bound Clr-DNA complex crystal structures.

**TABLE S6** Strains and plasmids, EMSA DNA fragments, Mutated DNA fragments, oligonucleotides.

